# Mouse Bio-behavioral Phenotyping Using a Digital Homecage Framework for Long-timescale, High-resolution, and Multi-factor Data Collection and Analytics

**DOI:** 10.1101/2025.04.25.650637

**Authors:** Nicolette Ognjanovski, Anjesh Ghimire, Pho J. Hale, David S. Kim, Ivo H. Cerda, Ethan Goldiez, Simeone Marino, Antwan Green, Paul J. Fitzgerald, Priya Vijayakumar, Deniz Kirca, Matthew Tong, Noah Muscat, Mitchell Cook, Kennedy Knopf, Mingyi Tang, Yang Chen, Lezio S. Bueno-Junior, Ridge Weston, Tangyu Liu, Jeremiah P. Hartner, Ivo D. Dinov, Brendon O. Watson

## Abstract

MOTIVATION

Long-term monitoring of behavioral and physiological processes is critical for understanding complex brain-based phenomena and disorders that develop over extended periods, such as chronic stress, circadian disorders, and metabolic conditions. While digital phenotyping is available in humans using smart devices, there remains a deficit in rodent models. To address this lack of long-term rodent phenotyping, we introduce the “Digital Homecage” (DHC) system. Our system uses accessible components and is designed for seamless integration with brain recording technologies.

The Digital Homecage (DHC) allows uninterrupted, long-timescale recording of more than 20 behavioral metrics in single-housed mice, captured at sub-second resolution via video, operant interactions, and wheel-running data. This report demonstrates the DHC’s capacity to enable continuous, automated tracking of behaviors like actigraphy, sleep, grooming, and food choice options over weeks, thereby opening up new avenues for longitudinal analyses of chronic conditions. Data collected reveal circadian patterns in multiple spontaneous behaviors, aligning with known nocturnal tendencies. The system’s potential to facilitate groundbreaking observations on the long-term behavioral correlates of various neuropsychiatric syndromes is aided by open-source software and relatively low cost. The DHC sets the stage for community-driven innovation, potentially transforming our approach to studying complex traits of brain function and behavior in laboratory settings.

## INTRODUCTION

Rodent neuroscience tools hold promise for studying brain-based phenomena and disorders, but many such phenomena develop over long periods or require long-duration monitoring to capture ^1–4^. Such slow-developing conditions include chronic stress, circadian disorders, aging, metabolic disorders, and others ^5–8^. Unfortunately, many such conditions have behavioral phenotypes that are not fully characterized by the standard short-term behavioral tests used over recent decades. Furthermore, when some of those standard tests are given repeatedly to delineate long-term trajectories of brain status, they often show changing behavioral results due to acclimatization, potentiation, stress, or other factors. On the other hand, measures of spontaneous behavior in the homecage, if done non-invasively, can be performed indefinitely for the lifetime of the animal and can reveal long timescale dynamics.

In an approach called “digital phenotyping” researchers use digital devices such as smartphones or wrist-based trackers to gather long timescale behavior and physiology data in humans, often over weeks or months. Using this longitudinal approach, scientists have discovered novel associations between behaviors and mental states ^9–14^. In the animal laboratory setting, scientists should be positioned to gather the same type of longitudinal daily activity data about rodents, since they live in animal facilities over weeks and months. However, behavioral testing typically occurs for only a few minutes or hours per week. Thus, rodent neuroscientists stand to gain huge amounts of new information by recording rodent homecage behavior 24-hours per day. Indeed, many aspects of behavior can often be better measured in the homecage than in the confines of arbitrary experimenter-defined assays ^15^.

To automatically capture such 24-hour rodent data we set out to create “rodent digital phenotyping”. While some homecage-based systems already exist, they often are not optimized for assessment of broad ranges of behavior or are expensive ^16^. Such systems have often also been wholly commercial in nature; whereas computer control systems, 3D printing techniques, and open science approaches have opened up many new lanes of innovation to individual labs such that they are no longer confined to commercial products. We have also reached a time in computing history where the kind of “big data” gathered from long-term recordings of rodents are relatively easily stored and processed. Finally, machine learning techniques can be brought in to complement more traditional analytics with such datasets.

The “Digital Homecage” (DHC) framework introduced here uses a combination of videographic, operant, and wheel-running data to capture 20+ behavioral metrics from a single-housed mouse at sub-second resolution, without interruption, over weeks. Broadly, this system can measure sleep, actigraphy, voluntary exercise, grooming, and exploration, as well as various consummatory behaviors. These are coupled with physiological assessment throughout the course of the experiment, providing the potential for detailed longitudinal analyses. This automated system will allow major new observations that define long-term behavioral correlates of chronic conditions and is made freely available via GitHub.

## RESULTS

### Digital behavior phenotyping system overview

We aimed to develop a platform capable of automatically registering a wide range of spontaneous murine behaviors in a standard laboratory homecage setting for weeks or months at a time. We prioritized affordability and accessibility including use of a standard rodent strain relatively low cost, simple design, standard parts, and compatibility with anticipated future, long-term brain recordings or manipulations. For those reasons, we chose to record C57BL/6J mice (Jackson Labs, JAX:000664) using a standard manufacturer homecage (Allentown) with simple modifications, surrounded by a system of operant feeders and water dispensers, camera, and a running wheel inside the homecage (Figure 1). Further detail for all procedures and designs is found in the methods section.

**Figure 1.**
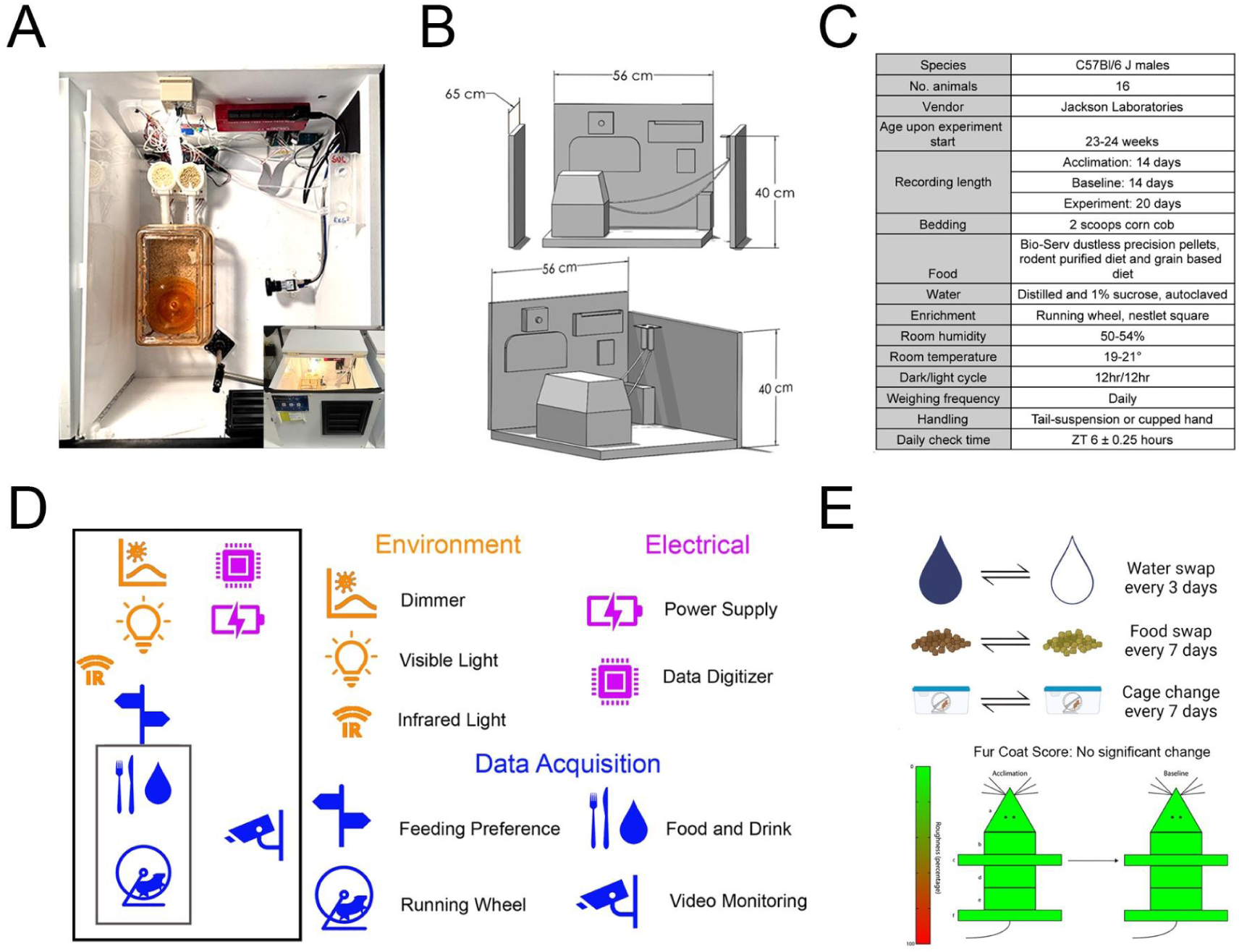
The Digital Homecage. **(A)** The DHC is a system designed to monitor the behavior of mice living in standard laboratory cages. Overhead view of a fully operational DHC including all components. Inset: Front view of DHC with lid opened. **(B)** Front and left side view of a 3D-representation of the DHC with top and front faces of the enclosing box removed. Dimensions listed for PVC sidewalls are provided in cm and cage and electronics are to scale. **(C)** Table showing experimental details and the environmental parameters controlled for in and around the DHC enclosure. **(D)** Pictographic example of the 10 main components of the DHC – elements shown are roughly aligned with DHC layout in (A). Environmental controls are shown in orange, electrical controls in pink, and all data reported in the results are shown as blue. **(E)** Top: regular events include water and food swaps every 3 and 7 days, respectively. Cages are changed every 7 days with food placement swap. Bottom: fur coat state was scored on water swap days in 6 different body regions (head, neck, fore paws, back, rump, and hind paws). There was no discernable difference between the first 2 weeks of recording (acclimation) and the second 2 weeks (baseline).

We designed and built 16 fully operational DHC units (Figure 1A), capable of chronically recording daily spontaneous behavior. The box is made of polyvinyl chloride (PVC) foam boards and is light-tight when closed, with a hinged top door to allow easy access for daily welfare checks and maintenance (Figure 1A, inset). Standard Allentown cages were placed within the DHC (Figure 1B) where they were surrounded by monitoring and feeding devices. To create a standard circadian light cycle, we diffused a constant 300 lux warm yellow LED strip through a sheet of translucent white plexiglass, simulating natural daylight conditions on a 12 hour:12-hour light: dark (LD) cycle (00:00 UTC is ZT0, independent of room light outside the light-tight DHC) with maintenance occurring during the 12-hour daylight period. The LED cycle is controlled by a LabJack T7 multifunction DAQ device (Figure S1, see Star Methods for further details). To enable 24-hour video recording of behavior during the dark periods, we used infrared illumination outside the mouse’s visually perceivable range placed behind the plexiglass diffuser. A filter was placed on the camera that permitted only infrared light to enter the lens, and the infrared LEDs remain illuminated continuously (during both the light and dark cycles) resulting in a uniformly lit backdrop against which the mouse’s activity can be seen with high contrast. The biological and environmental variables controlled within this DHC experiment are shown in the table in Figure 1C.

In total, there are 10 primary data streams continuously and automatically recorded within each self-controlled DHC at sub-second resolution (Figure 1D). The camera records continuously at 20Hz and generates video files that are later processed with software for video-based actigraphy analysis. This automated, high-throughput behavior recognition pipeline is capable of identifying multitudes of different behaviors at sub-second resolution. Non-video measures include nose pokes and beambreaks for two operant feeders and two operant water dispensers, and the rotary position of a running-wheel to gather exercise-related data. These digital signals are processed by the same LabJack T7 that controls the lights, which is connected to a recording computer that catalogs their values with sub-microsecond precision to disk. These feeding, drinking, and running data are recorded at 16hz (recent software rewrites now allow 100+Hz). All measures including camera frames are synchronized to the computer’s real-time clock, which in turn is synchronized with the NIST atomic clock at millisecond resolution and written into the resultant CSV files and video filenames as POSIX timestamps (aka UNIX time: a single number giving absolute number of seconds since January 1, 1970). In this way, all events are placed into a universal time frame for synchronization and later circadian analysis. These data streams are duplicated, backed up, and independently stored on local servers to be further processed offline (Figure S2). Data collected by the DHCs are supplemented by experimenter-executed daily software and animal welfare checks including weight, food and water refills, and intermittent fur coat score and cage changes ^17^. These data are recorded online via Google Forms.

We chose to focus on maximal behavioral detail from each individual animal. Therefore, we chose to single-house the mice; otherwise we would only have cohort-level accuracy. Given the added stressor of single housing, we closely monitored stress signatures via daily checks ^18^. Fur coat score (Figure 1E, bottom) was monitored every 3 days, coincident with water change days. As both overgrooming or under-grooming ^1^ create visible markers of animal distress ^7^, we also compared fur coat score during the first 2 weeks of acclimation relative to the following 2 weeks of baseline. This allows us to determine whether extended single housing contributed to any grooming abnormalities indicative of animal stress and welfare decline. We observed no statistical changes in fur coat score at any of the 6 positions assessed across the 28 days (p=0.4184, ANOVA for the effect of time, n=1422 across all body segments). Given their prior correlations with stress, measures of sucrose water preference (versus regular water) and fatty food preference (versus regular chow) were integrated into the DHC ^19–21^. These were carried out by putting sucrose versus regular water into our two water dispensers and regular versus fatty food in our two food dispensers. To control for and measure any location preference for the food or water dispensers, positions of sucrose versus regular water were swapped every 3 days and fatty versus regular food swapped every 7 days (Figure 1E, top). Cages were exchanged for clean cages on food location swap days.

Details of our control and acquisition system are stated in the Star Methods section (including all part numbers used). Using our 16 DHCs, we have recorded activity of 16 mice at a time, continuously for 8 weeks. We have not found specific temporal limitations on duration of recording but here we will present initial month-long data. In the next sections we will detail the technical components of the DHC before measuring these month-long baseline behavioral characteristics.

### Data acquisition components

In the experiment presented here, 16 wildtype male mice were recorded initially for 4 weeks (28 days). This provided adequate time for animals to normalize behavior to the novel environment. Since all data are stored with POSIX timestamps using the computer clock, repeated daily events can be analyzed relative to each other based on clock time or lights-on zeitgeber at 10:00 UTC (ZT0). In the rest of this publication, these 28 days are broken down into 2 weeks “acclimation” followed by 2 weeks “baseline” (as annotated in Figure 2 and throughout this text). Our completed DHC records three main data streams: operant food and water choice, running wheel use, and 24/7 video. While other automated animal monitoring systems have presented similar analysis for activity ^15,22–24^, the DHC records these three components concurrently at high resolution (Figure 2).

**Figure 2.**
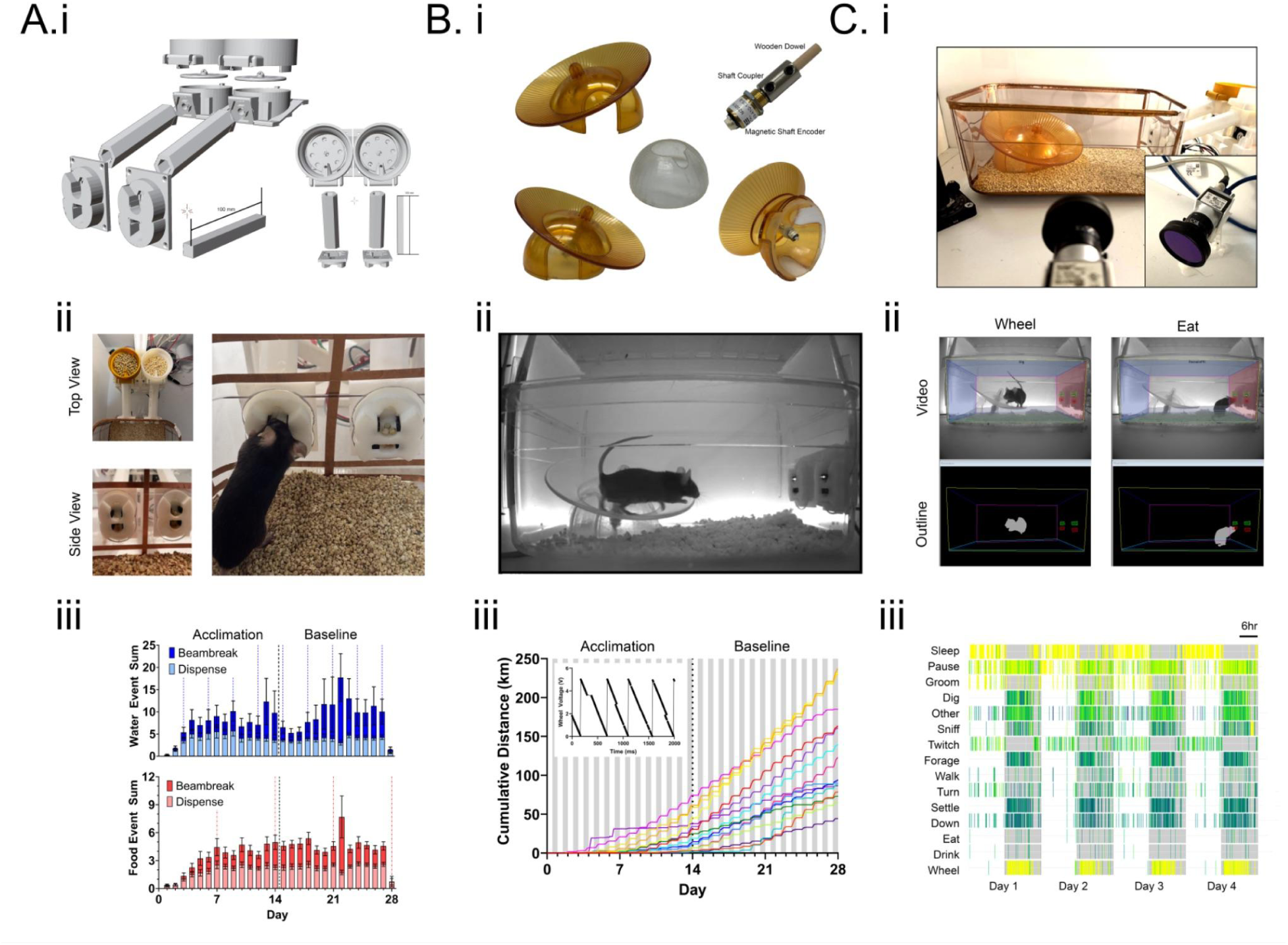
DHC main measurements. **(Ai)** For food and water dispensing, a 3D-printed structure fitted with two stepper motors houses regular and fatty food pellets. On each side, left and right, there is a food and water dispenser, but food and water types are randomized to left versus right for each animal and then are swapped every 3 or 7 days, respectively. **(Aii)** Images of food and water feeders (left) and a mouse interacting with the dispensing system (right). **(Aiii)** Counts over time for all water (blue) and food (red) dispense events (lighter color) and nose poke beam-breaks (darker color). This shows more beam breaks than dispense events for both water (p<0.0001, paired t-test) and food (p<0.0001, paired t-test). Vertical dashed blue lines indicate water swap days and vertical dashed red lines indicate food swap days. Data are shown across the acclimation (days 1-14) and baseline epochs (days 15-28). Data are averaged per day across all animals (n=16 mice, error bars are ±SEM). **(Bi)** Shaft encoder is fitted into the Fast Trac, commercially available running wheel, with 3D printed base. **(Bii)** Video frame showing positioning of the running wheel in the mouse cage. **(Biii)** Plot depicting cumulative distance travelled over time for each mouse in 1-hour bins (mice shown as individual colors). Shaded regions depict lights-off periods. Inset: absolute shaft position by generating an analog voltage output ranging from 0-5V. **(Ci)** An infrared camera recording 20 frames per second at 1.3 MP resolution continuously records activity inside the home cage from a side mounted position. **(Cii)** Batches of video files from the same cage are analyzed with HomeCageScan. **(Ciii)** HCS-derived raster plot for the top 15 behaviors observed + wheel for a representative animal across 4 days. Colors represent frequency of event per minute bin.

For food and water monitoring, we allowed mice the choice between 1% sucrose water versus regular water and fatty food (15%) versus regular food. To enable this, commercial plastic mouse cages had holes to accommodate two food dispensing modules and two water dispensing modules. The dispenser apparatus is a modification of on an open-source 3D-printed structure (https://open-ephys.atlassian.net/wiki/spaces/OEW/pages/79069188/Food+Pellet+Dispenser) and each is composed of the food dispenser in the original design plus a water dispenser at the mouse-facing port as well (Figure 2Ai). There are four beam-breaks, one for each of two food choices and two water choices, and each animal can see both left and right choices at all times. When the mouse breaks the photointerruptor beam with a nose poke, the corresponding food or water is dispensed (Figure 2Aii, right). The dispensed food or water is released immediately following the nose poke for it. Food is dispensed via a stepper motor mounted on a dispensing structure and water is dispensed by opening of a solenoid (Figure S3, see Star Methods for further details). Assignment of water and food type at each port was randomized for water type-food type pairing and right vs left dispensing unit. Mice had continual access to all food (blue, shown as a total of fatty and regular) and water (red, shown as a total of sucrose and regular) across the duration of the experimental timeline (Figure 2Aiii). We found that some mice repeatedly nose poke for food or water without consuming what is dispensed following the poke and even creating cage flooding. Therefore, we set a 3-second minimum time between dispenses (this timeout period is user-configurable). When measuring around-the-clock data, we find that dispense events are approximately 2/3 as frequent as nose-pokes, indicating that mice often poke multiple times within the timeout.

To track running behavior, a common commercial rodent running wheel (Bioserv, Mouse Igloo) is modified with a transparent 3D-printed dome to prevent mice from crawling under the wheel, thereby protecting the 3-pin micro connector carrying the analog signals to the LabJack device. This prevention of crawling under the wheel can also protect cables or tethers in future electrophysiologic or optogenetic experiments. The wheel’s position is encoded via a rotary encoder using 0 to 5V to denote radial 0 to 360-degree wheel position (Figure 2Bi). Voltage from the rotary encoder is fed into the LabJack that writes a timestamped line in the .csv when a voltage change is registered – therefore each voltage change denotes angular change of the wheel such as from running (Figure 2Bii). Each full rotation of the wheel (5V) corresponds to the mouse running 0.38 meters. Our findings show that across all animals (n=16), it takes mice ±7 days to begin continuous, daily use of the running wheel as seen by cumulative distance travelled (Figure 2Biii).

We use infrared lighting and an infrared camera filter to allow consistent quality 24-hour video acquisition despite 12/12 visible light cycling. A translucent acrylic panel diffuses a perpetually lit infrared backlight undetectable by mice to evenly illuminate the camera without interfering with the natural circadian cycle. We gather video data at 20Hz using a Basler camera (Figure 2Ci) controlled by Bandicam or StreamPix software. Video is triggered by a modifiable external timer to start and stop at intervals – for example every hour. Video data is saved to disk in real-time and processed post hoc using HomeCageScan (HCS, CleverSys) on a separate server (Figure S1C). We chose side-on camera orientation (Figure 2Ci-ii) for two reasons: 1) to minimize interference of video by cranial cables or tethers in future designs and 2) to be compatible with HCS. In Figure 2Cii we show both raw IR video and extracted animal outline for running and interaction with the dispenser representing eating or drinking. Figure 2Ciii displays an example of the top 15 major behaviors characterized by HCS as well as LabJack wheel data for four days (shown for a representative animal). These behaviors, for example ‘turn’, are summations of other more nuanced behaviors that fall under that classification, such as ‘turn right’ or ‘turn left’. We clearly observe consistent time-of-day (light vs. dark) effects on behaviors that are coincident with published reports of nocturnal behavior ^25,26^. Of note, these videos can also by analyzed by an open source software, LabGym, which can directly recognize behaviors without key points but requires users to train a library of behaviors to be recognized ^27^. To clarify, we have trained several behaviors in LabGym with good success but have not had time to train a full library. We may transition to LabGym in the future or combine it with HCS using each to best recognize specific behaviors.

### Multi-timescale analysis

Outputs of all DHC digital data streams can be analyzed at time scales ranging from seconds up to weeks. The ability to plot on different timescales allows for in-depth exploration of the slow-progressing responses that may result from environmental stress over time or for finding temporal interdependencies between behaviors. We assessed the three main data streams from the DHC (as shown in Figure 2), food and water consumption, running, and video analysis in bins of weeks, days, and hours with emphasis on 24-hour circadian timepoints across the day-night cycle (Figure 3).

**Figure 3.**
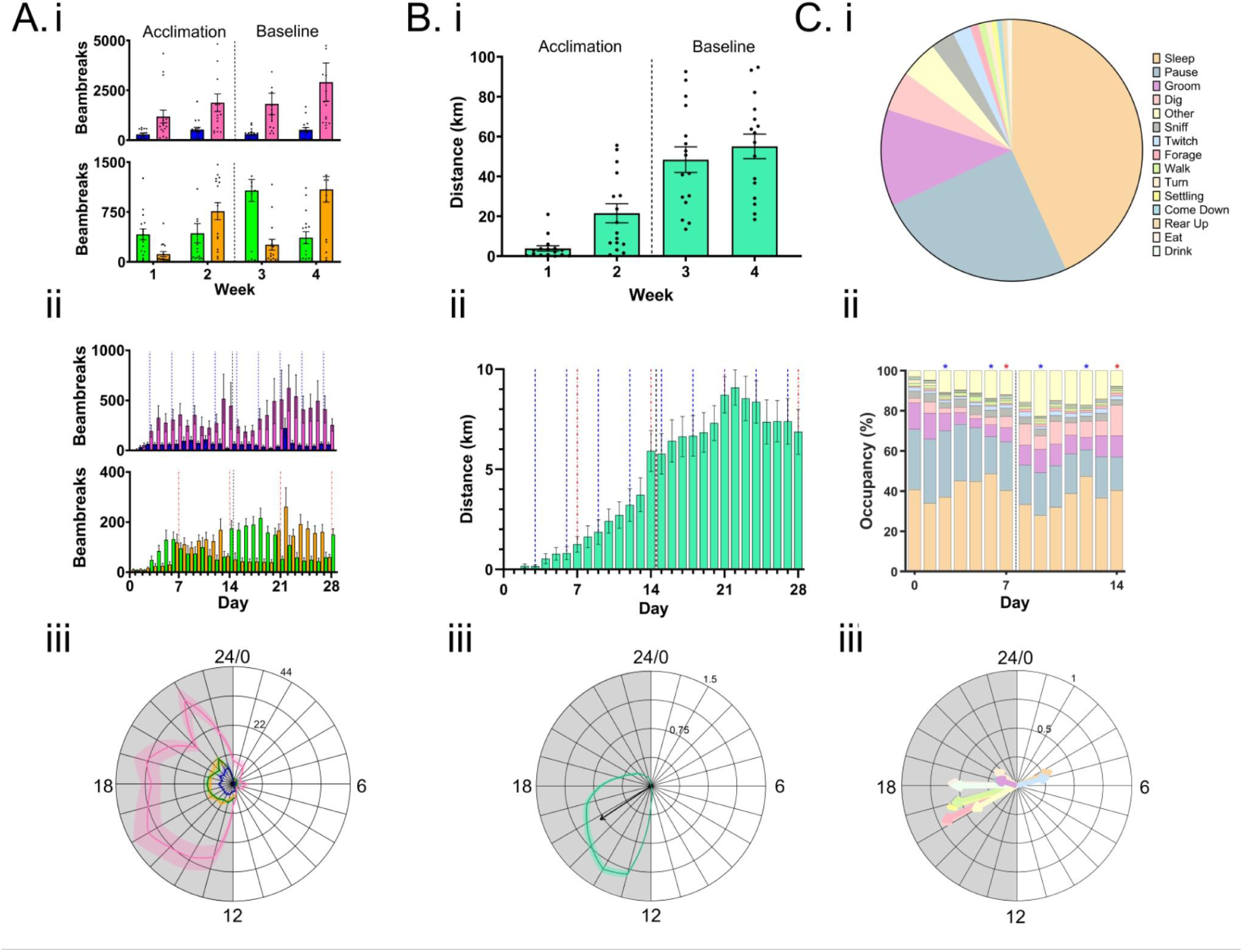
Weekly, daily, and hourly temporal patterns of measured behaviors. Time-binned examples of feeding/drinking as shown by beambreaks **(A)**, running **(B)**, and video **(C**) analysis across weeks (**i**), days (**ii),** and 24-hours **(iii)**. **(Ai)** Weekly binned data for beam breaks for water (top: blue-regular; pink-sucrose) and food (bottom: green-regular; orange-fatty). All bars are average across all animals for 7 days within each week. Error bars are ±SEM and all data points from each animal are noted as black dots. On average 13.93 ±1.14 pokes per hour for sucrose water, 2.78 ±0.31 for regular water, 4.08 ±0.45 for regular food, and 3.91 ±0.53 for fatty food. **(Aii)** Daily averages for food and water, with colors as in (Ai) allowing for visualization of animal preferences for positions in food (bottom panel). Bars show average across all animals for each day of recording. **(Aiii)** Hourly radial plot showing circadian time-of-day averages across all animals (dark line) ±SEM (lighter shading) for all presented food and water choices. Background is shaded to reflect day-night light cycle within the DHC. **(Bi)** Weekly running wheel distance for all animals. Displayed as shown in Ai. **(Bii)** Daily total distance run averaged across animals (7.39 ±0.24 km). Running distance stabilized at week 3 (p<0.001 Paired T test for first 14 days compared to last 14 days). **(Biii)** Hourly radial plot showing average km run per hour across all animals. Thin black line is average, green shading represents ±SEM with black vector indicating average circadian peak for running time and distance (ZT15.89). **(Ci)** Pie chart showing total experimental percentage of identified HCS behaviors for all animals. **(Cii)** Daily occupancy in each of the behavioral classes characterized as in (Ci) shown as a bar graph, colors as in Bi. Example is for an individual animal with asterisks indicating food (red) and water (blue) changes, with space in between bars after day 7 indicating cage change. **(Ciii)** Hourly radial plot of mean vectors for each behavior, colored as in Ci.

For food and water consumption, we analyzed the beambreaks (to represent the behavioral action of the mouse as opposed to relying on proper dispense functioning) to assess the *ad lib* demands for 1% sucrose (pink) and regular (blue) water as well as *ad lib* regular (green) and fatty (orange) food consumption (Figure 3A). First, we plotted the weekly timescale (Figure 3Ai) with the per-week average of normalized nose poke requests shown for all animals. The vertical dotted line shows the difference between the two weeks of acclimation and 2 weeks of baseline for all plots. Mice in general nose poked for sucrose water 3x as much as they sought out other food or drink options (13.93±1.14 pokes per hour for sucrose water compared to regular water [2.78±0.31], regular food [4.08±0.45], and fatty food [3.91±0.53], p<0.001, Browne-Forsythe ANOVA, Dunnett’s T3 multiple comparison for sucrose vs all others). Fatty food was not requested or dispensed significantly more than regular food (p>0.999, Browne-Forsythe ANOVA, Dunnett’s T3 multiple comparison). We next plotted on a daily timescale, allowing us to account for monitoring, food/water swaps, and cage changes as well as to improve temporal resolution (Figure 3Aii). Milestones of this particular experimental paradigm are shown with dashed vertical lines in each graph: water swap days are shown by blue lines and food swap days by dashed red lines. Animals show overall preference for sucrose water over regular water (Figure 2Ai) but seem to exhibit a place-preference in the positioning of their food, with clear swapping of fatty vs regular food eating coincident with food position swap days. This will be further analyzed in the next section. We lastly characterized hourly feeding and drinking around the 24-hour cycle to show circadian patterns. Mice, being nocturnal, eat and drink almost exclusively at night, with most dispenses occurring during the dark phase across all food and drink options (Figure 3Aiii, regular water: ZT16.97 ±0.34, sucrose water: ZT18.29 ±0.16, regular food: ZT17.64 ±0.15, fatty food: ZT17.74 ±0.17)

We then characterized weekly, daily, and circadian wheel running data (Figure 3Bi-iii). Weekly averages across all animals for the 2 weeks of acclimation and 2 weeks of baseline were plotted (Figure 3Bi) and show a majority of animals (n=12) level off of running behavior after the 2-week acclimation period running ∼7 km/day (Figure 3Bii). On average the mice run 1.805 km/day during acclimation vs. 7.39 km/day average during baseline days (p<0.001, paired t-test for first 14 days compared to last 14 days), which is consistent with free-running behaviors in populations of wild mice ^23,24^. This suggests that under laboratory constraints, with any possible stress due to single-housing, mice do not behave naturalistically until 14+ days of consistent exposure to the running wheel. We next analyzed data within repeated 24-hour daily cycles, in hourly bins, and averaged across animals to create circadian plots of daily running rhythms (Figure 3Biii). The average distance run within each hour shows that this cohort of animals ran mostly at the beginning of the dark period (Figure 3Biii, green line) which is the expected active phase for the nocturnal mouse. We quantified the mean vector for this running activity for all recorded animals (Figure 3Biii, black arrow) and find maximum activity around ZT15.67 ±0.08. This may coincide with bursts of activity at the start of the active phase after 12 hours of increased sleep.

For video analysis, following HCS software-based scoring of multiple behaviors from video, we summated across the two weeks of baseline recording and generated a pie-chart displaying percentage of total video frames spent in each behavior across all (n=16) animals for all days for which we ran HCS (Figure 3Ci). After grouping the 44+ behavioral classifications provided by HCS as described previously, we found that the top 15 behaviors account for 92.76 ± 0.43% of all time recorded with 10 additional behaviors such as chew, circle, hang, jump, land, and stretching occurring intermittently with less frequency and are therefore not analyzed in this publication. We quantified the top 15 behaviors in daily bins, with bar graph portions indicating total time spent in each behavior across the day (Figure 3Cii, shown for a representative animal). Sleep, as characterized by HCS accounted for 43.19% of average daily behavior, the largest behavior across all days of HCS analysis. Across these 14 days, the percent of time varies slightly on a weekly cycle and sometimes around water swap day (Figure 3Cii and S4). These results reflect an expected amount of inter-animal individuality.

In order to possibly account for this, we further characterized circadian phase of behaviors across all animals using a mean resultant vector of the hourly occupancy for each behavior (Figure 3Ciii, colors same as in Ci). While most behaviors occurred during the dark (active) phase, the “sleep” and “twitch” behaviors (Figure 3Ciii, orange and periwinkle arrows, respectively) occurred during the light (inactive) phase (ZT4.50 ±0.22 and ZT4.99 ±0.24, respectively for sleep and twitch). This is of note as twitches are often associated with REM sleep or sleep more generally ^28–31^. This preference of sleep-based behaviors in the inactive phase then serves as initial validation of the video scoring methodology. Furthermore, the phase (ZT18.12 ±0.22) of the “eat” behavior via video scoring was similar to the phase of food consumption using our operant feeder data (as shown in Figure 3Aiii ZT17.69 hours average eating time across both food types). HCS calculated average drinking occurs at 17.98 ±0.22, which is again similar to the water beambreak timing of about ZT17.5 hours. These time-of-day preferences validate the accuracy of the commercial automated behavior scoring algorithm relative to hardware tracking of these actions.

### Preferences in animal feeding behaviors

Mice show preference for sucrose water vs. regular water (Figure 3A) despite water location swaps, but weekly food location swaps correlated with a bigger swing in fatty versus regular food consumption (Figure 3Aii). Therefore, we further quantified the tendencies of animals to nose poke at specific locations of dispensers. Beambreaks for both water and food types were summed for each day of the experiment across all animals (Figure 4A-B, respectively). We observed swapping peaks for each position, in alignment with mice following the sucrose water dispenser position (Figure 4A) as well as noting general preferred interaction with food at position 2 (Figure 4B). Across the 4 weeks of acclimation and baseline, the total amount of water, independent of position remains stable (5.28mL/day, average 2.634 ±0.205 mL for position 1 and 2.648 ±0.226 mL for position 2, p= 0.9508, paired t-test for entirety of recording), data not shown as there was no equal measurement in terms of number of pellets monitored for food. During our 4-week recording time, mice lost weight across the first 3 days, possibly in line with familiarizing themselves to the feeding setup and running wheel. Generally, weight rebounded starting on day 4 of acclimation and stabilized into the baseline period (Figure 4C, 29.95g ±0.137g, p= 0.0624 Welch’s ANOVA test for average weight over time). Blue and red lines indicate dates of water and food location switch, which did not affect weight over the 4-week recording.

**Figure 4.**
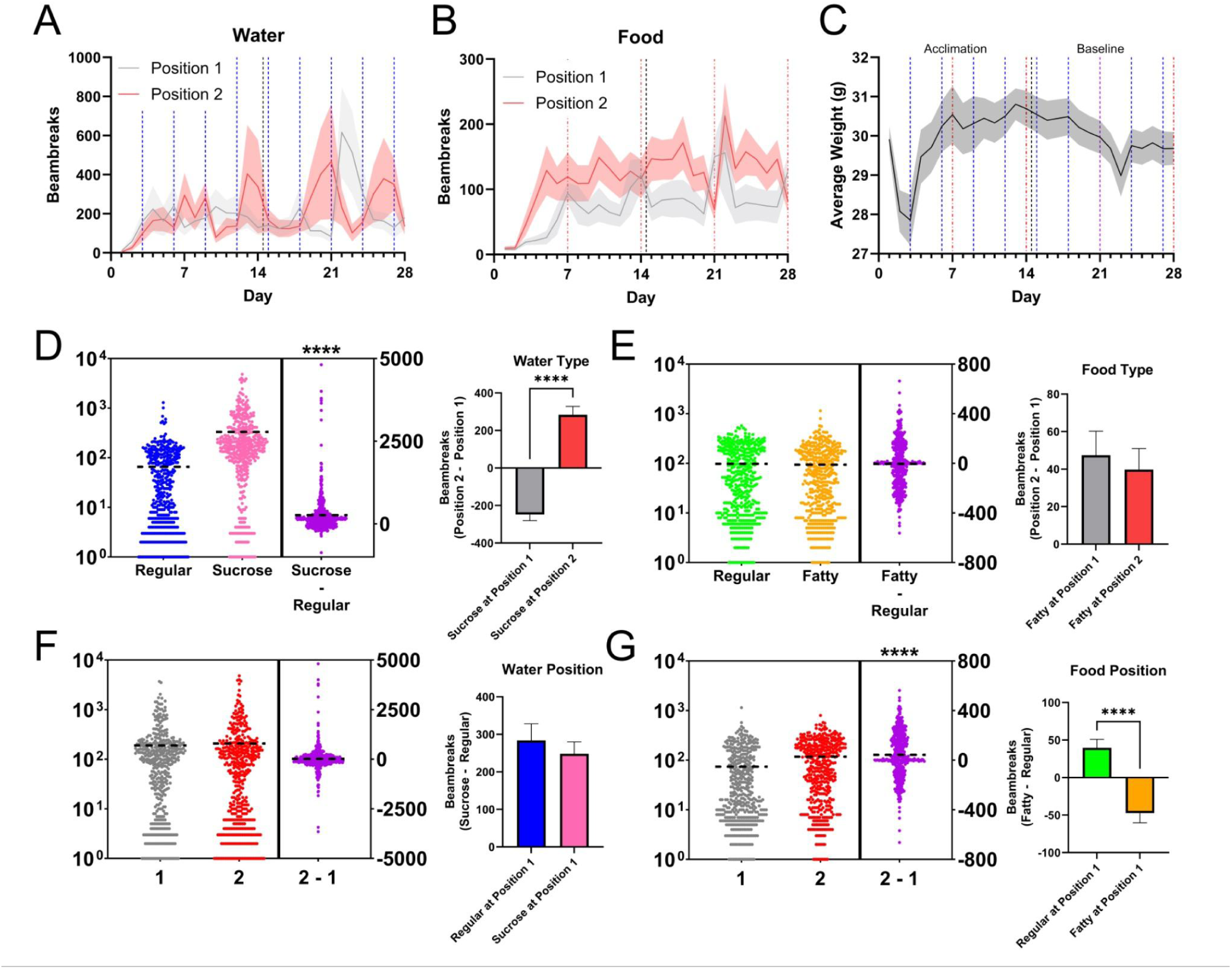
Given operant choice, mice have water sucrose preference but food position preference. **(A)** Total beambreaks of all water at position 1 (grey, closer to power supply) and position 2 (red, closer to diffuser) did not statistically vary across the recording time period. Blue vertical lines show water swap days. Black line is cage change and food swap. **(B)** Same as in (A) but for food beam breaks. **(C)** Daily weight was averaged across all animals and showed leveling off over the course of the experiment after initial 4-day adjustment. Vertical lines as indicated in (A-B). **(D**) Left: Regular water (blue) and 1% sucrose water (pink), are plotted as total beambreaks across all animals for all recording days. Each point is a daily average for one animal (448 data points total) with black horizontal lines showing overall average. Difference between sucrose and regular water shown in purple, with preference for sucrose (****p<0.0001, Paired t-test). Right: The difference between sucrose beam breaks vs regular beam breaks at position 1 (grey) vs. position 2 (red), show the sucrose preference occurred independently of position, (****p< 0.0001, Mann-Whitney test). **(E)** Regular food (green) and fatty food (orange) are plotted as total food beambreaks with difference shown in purple, with no preference shown (p=0.6618 paired t-test). The lack of food preference was not affected by positioning when normalized for overall preference of fatty vs regular food (p= 0.4785 Mann-Whitney test for position 1 [grey] vs. position 2 [red]). **(F)** Comparing all water beambreaks for position 1 (grey), position 2 (red), and difference (purple) showed no preference for water placement (p=0.5585, paired t-test). When position is compared between regular (blue) and sucrose (pink) water there was no preference (p=0.6130 Mann-Whitney test). **(G)** Comparing preference for food showed overall preference for position 2 (red) compared to position 1 (grey), independent of type of food (****p<0.001, Paired t-test). When position preference is measured between regular (green) and fatty (pink) food, there was a preference for regular food at position 1 and for position 2 for fatty food (****p<0.0001 Mann-Whitney test).

To account for swapping the position of water every 3 days, we quantified food and water nose pokes in position 1 (closer to the power source) versus position 2 (closer to the diffuser panel and the IR light) regardless of which type of water was at each position. We also quantified by water/food type. First, we observed an overall preference for sucrose (p< 0.0001, paired t-test) across all animals (Figure 4D, left. blue=regular water and pink=sucrose, as in previous figures). This preference was strong enough that sucrose water was preferred regardless of dispenser position. This was shown by findings of significantly non-zero but opposing values for position 2 minus position 1 regardless of sucrose position – the opposing values showing that sucrose was more preferred than position (Figure 4D, right). Using related logic, sucrose minus regular water was positive regardless of sucrose position (Figure 4F, right), again indicating that sucrose preference overrides place preference for water.

There was no preference for food type, regular or fatty (Figure 4E, left, p=0.6618, paired t-test). The position preference in food data is shown similarly to type preference for water, via two parallel subtraction tests: position 2 minus position 1 and fatty minus regular food. In the test of position 2 minus position 1 for food data, we found a consistently positive value regardless of which position had fatty food (Figure 4E, right), indicating that position preference overrides type preference. Interestingly, a subtraction of fatty minus regular did depend on position of fatty food (Figure 4G, right, p< 0.0001), meaning position was more important than food type. The preferences noted represent the beginnings of multi-variate analysis we can do with data acquired with the DHC.

### Web-based data processing module

We have built the DHC framework as an integrated development environment (IDE) based on the statistical language R. This allows us to utilize sophisticated backend data analytics R libraries, statistical learning and AI tools, as well as RShiny frontend user and machine interfaces. All data are stored in a manner that is available to our RShiny server (Figure S1). Both video- and feeder/water/wheel-based data are initially saved in un-binned format with fully precise, sub-second timestamping. R code then bins data into days, hours, and minutes for presentation and analysis online (Figure 5). On the RShiny server, users are able to view data via a web browser interface for quality control or basic assessment. Feeding, drinking and running wheel data are saved as .csv files (one line per time point) and read by the server every 24 hours and can therefore be viewed freshly each day of the experiment. Video data require manually initiated setup and initiation of automated behavior scoring software but are then saved as a different .csv file on the same server. Therefore, video data viewing is determined by human pacing of video analysis.

**Figure 5.**
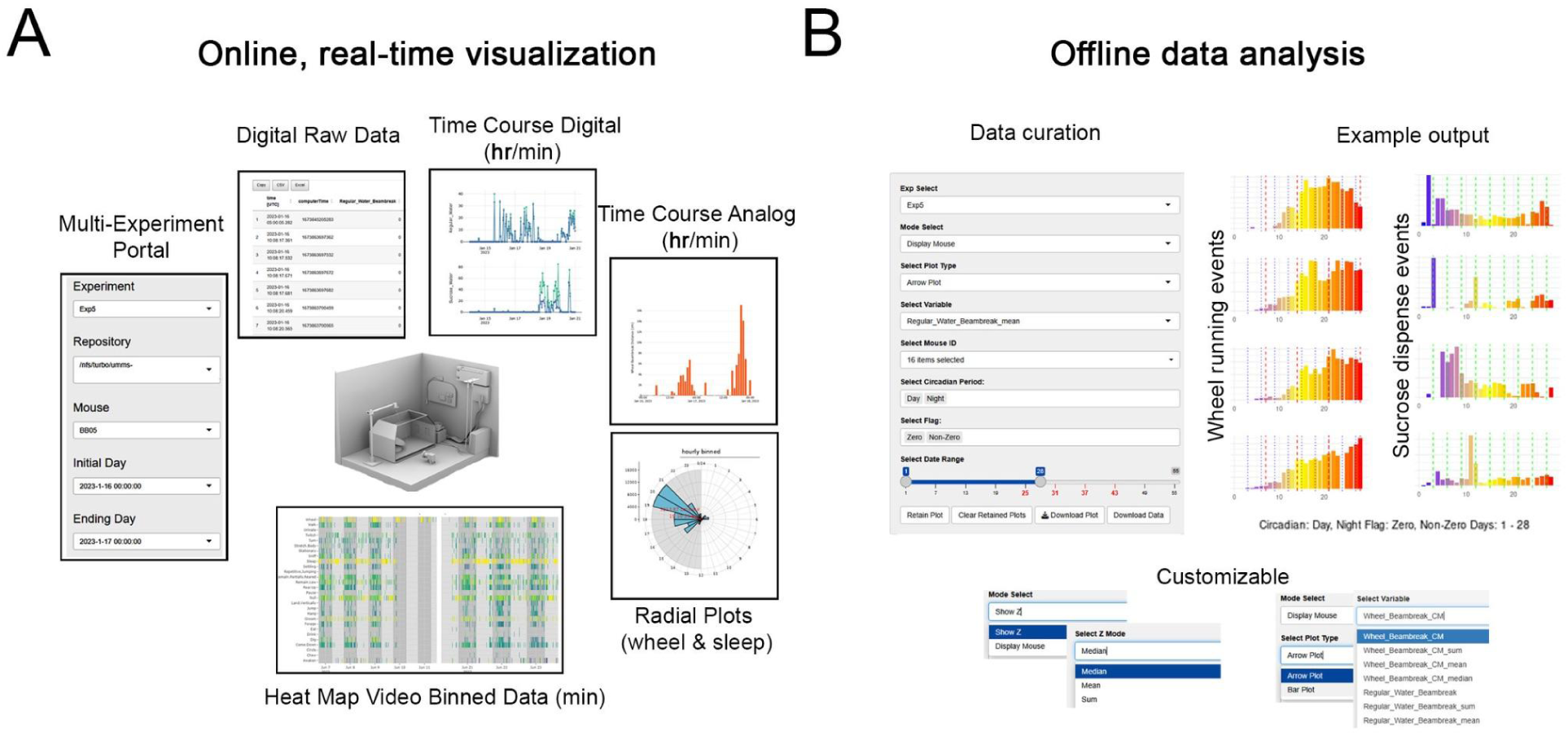
Data interfaces. The RShiny server allows users to view data via a web browser interface for quality control and data extraction. **(A)** Ingestion and aggregation of events into time bins is performed automatically every 24 hours and made accessible to our RShiny server. **(B)** A separate interface further extracts data for more detailed analysis and statistical testing implementing mean, standard deviation, and tracked sums is also available offline.

The server has two major modules: One focused on time-bin based displays (Figure 5A) and another with focus on statistics and population activity (Figure 5B). The RShiny app supports real-time interactive online visualization supporting quick explorations and identifications of potential failures in either the hardware or software stream. Data can be viewed at the individual animal level. Additionally, on completion of experiments, all backed up data on the RShiny server are available with statistical options (e.g., mean, median, and sum) data presentation customization (Figure 5B). This interface also allows export of both graphics and spreadsheet-format versions of data, binned at minute, hour, and day for further analysis using other software programs.

## DISCUSSION

The DHC system presented here enables detailed longitudinal recording of dozens of behaviors at sub-second temporal resolution for multiple animals at once via multiple DHCs. Thus, it offers both high temporal and behavioral resolution to allow long term, precise digital phenotyping. The data from this system can be analyzed at multiple timescales: from multi-day spans of interventions (i.e. blocks of stressors, or drugs), to days (behavioral paradigms), to hours (i.e. circadian assessments), and even to minutes, seconds, or finer to match with brain activity measures. The multi-dimensionality of the data can also allow different classes of questions to be asked ranging from metabolic (i.e. food intake, weight, exercise output) to circadian analyses, to intervention-based questions. Multi-behavior interactions over time or per animal can also be analyzed. In the future, brain-behavior linkages at a variety of timescales are also possible with the planned integration of *in vivo* brain recording systems into the DHC. This multi-dimensional and multi-level analysis can be of great utility in mechanistic studies of human diseases using behavioral phenotyping which can be bridged by the powerful neuroscientific tools available in rodent systems. Since our system is non-commercial and designs will be openly shared, we hope it will serve as a starting point for further refinement and development by those in the community.

We specifically focus on spontaneous behavior given the frequency and importance of spontaneous behavioral changes in neuropsychiatric syndromes in humans ^32,33^. Thus, this high-throughput and high-resolution approach can establish integrative biomarkers for disease models or other long-developing temporal processes such as stress, aging or others.

### Initial findings

We have been able to consistently use the DHC over multiple weeks in 16 mice simultaneously. Our data gathering and presentation platform allowed us to quantify data over a variety of timescales. We observed that mice show significant circadian modulation of many daily behaviors including actigraphy, sleep, eating, drinking, wheel running and foraging.

Importantly, we observed that wheel running requires approximately 14 days to reach a steady state, as well as feeding and drinking behaviors taking several days to normalize after experimenter interventions like water swap (place preference differences) and cage changes (video differences). This could be a matter of learning the wheel initially and then building the interest and habit, or as this is the animals’ first exposure to the wheel, it could be a factor of novelty, with mice being uncertain how to interact with the wheel. Or perhaps there is a delay for mice to become comfortable with the entire DHC and thus delaying naturalistic running tendencies. Recording of true steady state behavior is therefore enabled by our long-term monitoring and has important implications for future work, since many measures may be best performed when animals are at steady state, rather than at earlier or shorter term timepoints. The use of our system can enable just this kind of long-term observation, and our software interface can even allow users to track progress to steady state before initiating recordings that will be used in analyses.

We alternated food and water placements to assess the consistency of preference for regular vs fatty food and regular water vs sucrose water. We found that in general there is a preference for sucrose water over regular water, but no preference for fatty vs regular food. A majority of animals did prefer a specific food dispenser, possibly driven by the initial locations of each option. Preference for position may result from unobservable effects, e.g. the mouse sensing of IR light (position 2 is located closer to the diffuser panel). On the other hand, sucrose water was consistently preferred despite location swaps. These data suggest high inter-individual variability across such a long recording time that may be explained by other factors such as running or behavioral changes. The DHC also offers functionality to monitor weight, diet, and exercise, including the relative balance between each. We saw consistent weight over the last two weeks of the recordings, implying self-regulation of caloric status despite changing content including increased calories through sucrose water, ability to choose fatty versus regular food, and the new opportunity to exercise with a running wheel (not in their previous cages).

Additionally, circadian quantifications are efficient with this system with many behaviors showing modulation by time of day. For example, wheel running and sucrose water consumption were strongly modulated to occur most in the early dark hours. On the other hand, sleep and twitch happened mostly in the light hours in this nocturnal animal. Grooming was among the least circadian modulated behaviors, as it occurred relatively constantly in every hourly time bin (average 6.99 ± 0.134 minutes each hour) with no clear preference for day or night as observed in most other behaviors.

### Advantages of the DHC system

Many neuropsychiatric syndromes, and indeed most behavior in general, are based on self-motivated spontaneous drives which are often well-monitored at home. A homecage-based approach can thus behaviorally complement experimenter-defined assays; as a result, some researchers have studied more naturalistic circadian time-scale changes in the homecage ^22–24,34^. We are able to perform circadian analyses on all of our variables presented in this paper, which all are spontaneously performed in 12/12 light cycling.

The observation of multiple dimensions of free homecage behavior over days, weeks, or months has a number of advantages. First, it can quantify baseline behavioral attributes of individual animals, thus capturing inter-individual variation and possibly providing predictors of subsequent responses to interventions. Second, it can lead to data-driven multi-dimensional biomarkers - including specific alterations at particular timepoints – to stressors, drug treatments, gene silencing, or other interventions. This stands in particular contrast to traditional experimenter-initiated tests that do not capture natural behavior and are often carried out at arbitrary times. Third, it can also lead to biomarkers that are based on temporal evolution of multiple elements. For example, after chronic psychosocial stress sleep changes could precede alterations in eating or grooming. Such fine experimental controls may support studies examining deeper behavior, stress, biomarkers, and various conditions.

### Limitations of the study

Our system uses single-housed rodents, and single housing has been noted as a stressor ^35–37^. The decision to record single animals in this system was multifold. First, a long-term aim of our system is to link behavior to brain measures such as electrophysiology. In such experiments, multiple animals are infeasible since they often chew or destroy each other’s implants which can damage the implants or at least ruin behavioral phenotyping efforts due to their preoccupation. Second, the behavioral resolution is decreased with each animal added to the DHC environment. Presently, we do not yet have reliable means of differentially tracking conspecifics. Radio frequency identification (RFID) technology is one option that can give some resolution, but it requires surgery, and the spatial resolution is poor compared to a mouse cage – meaning frequent inability to distinguish between mice in any given location ^38,39^. Additionally, video analyses commit errors of swapping individual animals, even in neural network-based systems ^27,40–42^. Until future technical development occurs in the ability to reliably track single animals in a group setting, a single housing system provides significant advantages for many experiments. Importantly, stress has demonstrable effects on single housed animals, indicating that the stress of individual housing is not saturating ^17^. Running wheels such as those in our cages can reduce the measurable effects of stress in single-housed animals ^43,44^. An alternative could be to allow some hours per day for socialization with familiar conspecifics to attenuate stress effects, though this is not yet explored. Importantly, a recent study showed that even in single-housed animals, stress shows effects above and beyond controls ^17^.

Another point of difficulty is the high levels of inter-individual variability apparent in datasets with such high resolution. Essentially, this is a problem based on having more information than what might result from testing in more traditional manners. For example, Figure S4 shows individual animal video analysis complexity. We were initially concerned to see some of this variance, but despite this high variability, we see population-wise effects in good alignment with prior studies, such as sucrose preference and circadian modulations ^45^. Importantly, higher single-animal resolution should aid true phenotyping and generation of reliable biomarkers. Such variability can be considered to be an intrinsic feature between individuals, leading to even deeper understanding of behavior-environment conditions in human populations. Therefore, we are simply capturing and fully measuring variability that is already intrinsic to the animals, lending us more information for fuller understanding of populations.

### Future directions

The DHC may be applied more broadly after this initial development and characterization of the baseline recordings of spontaneous activity as a novel way of understanding mouse behavior. Behavioral tendencies, sex differences, or finding stereotyped behavioral sequences (e.g. nesting before sleep or drinking before eating) will all provide new fundamental data for mouse neuroscience ^46^. Additionally, sex differences may become apparent and can be quantified in spontaneous behavior. Beyond baseline homecage behavior in wildtype mice, behavioral questions can be examined using the DHC including drug and stress response, optogenetic or chemogenetic interventions in specific brain circuits, genetic mutant behavior changes, and others. Additionally, the DHC as presented is currently being scaled up to accommodate housing for rats (*Rattus norvegicus*).

There are potential scientific directions for the DHC we have only touched upon thus far including metabolism, diet, and exercise. With this system, researchers can examine impacts of mid-experiment changes in food or water choices, exercise availability or difficulty (increased resistance or availability at only certain times of day). Outcomes of these could be measured via weight, but also via fecal testing of microbiome or other content, or blood tests ^47,48^.

Circadian interventions represent yet another potential use of the DHC, with approaches to quantify single or multi-behavior patterns across the day. While the findings here report a baseline, they could be manipulated including by changing timing of presence of food, light, or exercise wheel. Outcomes can be measured behaviorally, in weight or with physiologic measures. Moreover, the DHC can be re-deployed for diurnal species such as Nile grass rats (*Arvicanthis niloticus*).

In the future, we intend to couple these behavioral measures with physiologic recordings including electromyography, electrocardiography, respiration monitoring, or body temperature monitoring. Such recordings can be synchronized to this sub-second resolution system to yield fine-scale brain correlates with specific spontaneous behaviors. Brain recordings can be at the electroencephalographic level, depth recordings with isolated units, fiber photometry, or calcium imaging using miniscopes. This is potentiated by the high temporal resolution and the side-view nature of our camera system which tolerates tethered animals particularly well. Additionally, brain or body control mechanisms can be added using electrical or optogenetic stimulation or drug pumps – including via closed loop approaches. Cognitive tests can also be integrated into the system using touch screens coupled with the dispensing of food or water.

Finally, the analytical pipeline can be further refined to automate profiling of individual animals, circadian profiles, patterns of sub-groups of animals or behavioral sequences using both traditional and machine learning approaches.

## Conclusion

In the DHC we have developed a new system for recording spontaneous homecage behavior of mice that live in our laboratories. This system can provide new insights into mouse behavior for a variety of biological questions and can be coupled with other recording methodologies or models for broad utility in neuroscience or other behavior-dependent rodent questions.

## RESOURCE AVAILABILITY

### Lead contact

Further information and requests for resources should be directed to and will be fulfilled by the lead contact, Brendon O. Watson.

### Materials availability

All data and analysis materials generated in this study have been deposited to Deep Blue Repository at the University of Michigan. Deep Blue Repository and Research Data Services (DBRRDS) provides support for the access to this information, which is open-access.

### Data and code availability

All code used in processing of these data is available on the GitHub as noted in the text and in Materials availability. Specifically, in subfolders ‘Software’, ‘Procedures’, and ‘Systems’ of the GitHub page. Any additional information required to reanalyze the data reported in this paper is available from the lead contact upon request.

## ACKNOWLEDGMENTS

This work was supported by the NIH (MH107662), the Pritzker Neuropsychiatric Research Consortium, The A. Alfred Taubman Medical Research Institute, The Eisenberg Depression Center, and the University of Michigan Neuroscience Scholars fund to BOW. Additional funding from the Michigan Medicine Sleep Disorders Center: Gilmore Sleep fellowship and Marie Curie Horizons 2020 Leading Fellowship (No.707404) to NO provided while at the Leiden University Medical Center. NSF (1916425) and NIH (T32GM141746) funding provided partial support for SM and IDD.

## AUTHOR CONTRIBUTIONS

Conceptualization, B.O.W.; Methodology, N.O., A.G., P.J.H., D.S.K., I.H.C, S.M., A.G., P.J.F., P.V., M.T., L.B., J.P.H., and B.O.W.; Software, P.J.H., D.S,K., S.M., P.V., D.K., N.M., M.T., Y.C., L.B. and T.L.; Formal Analysis, N.O., E.G., and K.K.; Investigation, N.O., A.G., P.J.H., D.S.K., E.G., A.G., P.V., D.K., M.T., N.M., M.C., K.K. and R.W.; Resources, A.G., P.J.H., D.S.K., I.H.C., S.M., B.O.W., Data Curation, N.O., A.G., E.G., S.M., A.G., and K.K.; Writing—original draft, N.O., A.J., I.H.C. and B.O.W.; Writing—review & editing, N.O., A.G., D.S.K, A.G., E.G., B.O.W. and P.J.H.; Visualization, N.O., E.G., S.M., A.G., K.K., and B.O.W.; Supervision, N.O., A.G., S.M., I.D.D., and B.O.W.; Project Administration, B.O.W.; Funding acquisition, N.O., H.A., I.D., and B.O.W.

## DECLARATION OF INTERESTS

The authors have no conflicts of interest to declare for this publication.

**DECLARATION OF GENERATIVE AI AND AI-ASSISTED TECHNOLOGIES**

During the preparation of this work, no generative AI or assisted technologies were used.

## SUPPLEMENTAL INFORMATION

Document S1. Figures S1–S4

Supplemental Figure 1. Electrical Components and flow of information and power between the different components of the DHC

Supplemental Figure 2. Data transfer, backup, and storage for processing

Supplemental Figure 3. Food and Water control in the DHC

Supplemental Figure 4. Daily behavioral breakdown of HCS video behavior

## EXPERIMENTAL MODEL AND STUDY PARTICIPANT DETAILS

All animal husbandry and surgical/experimental procedures were approved by the University of Michigan Institutional Animal Care and Use Committee (PRO00011423 - BOW). Male C57BL/6J mice (n = 16; IMSR_JAX:000664, Jackson) were individually housed in standard caging modified for the DHC (see details below). Mice were between 3-6 months of age for the duration of the experiment. Environmental light was maintained on a 12:12 light:dark cycle (lights on at Zeitgeber Time [ZT]0) with food and water available ad libitum. Prior to the start of each experiment, bedding and running wheels were added to each mouse cage in each DHC and the brightness of the visible LED was calibrated. Every 7 days, cages were replaced with clean cages with fresh bedding. Maintenance and care, including institutional permission and oversight information for the experimental animal use were overseen by University Laboratory for Animal Medicine (ULAM) at the University of Michigan with approval from IACUC.

## METHOD DETAILS

### Box Design

Exterior Encasement: A 68.6 cm x 55.9 cm x 43.2 cm box made with white ¾ inch Foam PVC (American Plastic Solutions) encloses the DHC system. The enclosure has a 25.4 cm x 25.4 cm (10” by 10”), light-tight louver vent in one side wall that maintains temperature, air composition, and humidity to an acceptable level, offsetting the heat produced by the various electronic components and light sources that the system deploys.

Mouse Cage: A standard commercial animal cage (Allentown, #PC75JCL) with standard bedding (Nestlets, LabSupply) provides the housing for our chronic measurements. Two 3.8 cm x 5.1 cm figure-eight shaped patches matching the shape of our two dispensing units were drilled out of the Arduino-facing side of the mouse cage (one of the smaller walls of the rectangular cage). A small 1.27 cm hole is taken from the bottom of the cage to introduce the 3-pin micro mating connector of the Running Wheel System. Mice are allowed to eat, drink, run, behave and sleep *ad libitum*. Prior to the start of each experiment, bedding and running wheels were added to each mouse cage in each DHC and the brightness of the visible LED was calibrated.

### Lighting and Schedule

We used a 12:12 light:dark cycle for these experiments, although the schedule can be changed via LabJack code. All visible LEDs were on daily at 05:00 UTC for 12 hours, and IR LEDs for camera illumination are on 24-hours daily, diffused through a #7328 Translucent White Acrylic Sheet. Visible light is emitted from 6 LEDs that output 6000 Kelvin light (to simulate bright daylight) at approximately 40 lux encased within a rectangular prism of white translucent acrylic that measures 0.635 cm x 34.3 cm x 61 cm). 40 lux brightness is set via Light Dimmer ZDM-01 from JKL Components and is measured at the center of each mouse cage with the mouse cage lid attached.

### LabJack

There is one computer for each DHC, connected to a camera and a LabJack T7 digitizer (with a CB15 Terminal). The LabJack timestamps events and continuously stores them in two .csv files on the DHC-specific computer to which it is connected by USB. Food and water beambreak and dispense data go to one file, while running wheel data go to a different .csv. For food and water, LabJack records as binary (digital) the TTL voltage outputs from the Arduino: half of channels recording beambreak times and half recording dispense commands to water and food motors/solenoids. LabJack records as analog (16-bit) the 0-5 V from the running wheel rotary encoder. All channels specified in Supplementary Figure 1B. For each event, line is appended to the end of a .csv which includes a POSIX time entry in the first column. Additionally, we employ the LabJack T7 5V DC output to precisely control the light cycle of the boxes as described in the Visible Light Relay Assembly subsection. The computer for each DHC also writes .mp4 video files from the camera for each DHC and stores them to a local RAID. All files are automatically backed up daily to distant cold storage.

### Electrical System Design

The Circuit Panel Assembly takes in 12V of power from the AC/DC converter (Mean Well USA Inc, EDR-75-12) and distributes it to key components of the DHC (as shown in Supplemental Figure 1) including the Light Relay Station, the Arduino, and the Motor Shield. The Circuit Panel Assembly consists of a 1) Power Distribution Board Assembly, 2) 12V Arduino Barrel Jack Wire Assembly, 3) Buck Converter Power Assembly, 4) 12V Visible Light Relay Power Wire Assembly.

1. Power Distribution Board: the right half of a red Breadboard Mini contains the system’s 12 VDC power rail, which is supplied directly by the high-capacity power supply which gets power from the wall. The Arduino, the infrared backlight, and the visible LEDs all operate directly from this 12V source. On the other hand, the Motor Shield Assembly also draws from this supply after it is stepped down to 9V-DC via a buck converter (LM2596, Valefod). The left half of the breadboard contains the independent Water Solenoid Valve Driving Circuit, which allows the Arduino Motor Shield to control the opening and closing of solenoid valves for water dispensing.
2. 12V Arduino Barrel Jack Wire Assembly. This assembly consists of a 53 cm-long cable connected to the Power Distribution Board on one end and a male barrel jack connector on the other end. It provides 12V DC power to an Arduino Mega on the Arduino-Motor Shield Construct.
3. Buck Converter Power Assembly. The buck converter steps down the 12V input from the Power Distribution Board to provide 9V of power to the Motor Shield. It consists of a 10 cm-long 2-conductor cable connecting the Power Distribution Board’s 12V and GND rails to the input terminal of a buck converter that steps down the voltage output to 9V. A 72 cm-log wire then connects the buck converter’s output to the Motor Shield Assembly, supplying 9V of power.
4. Visible Light Relay Assembly. Made of a 51 cm-long 2-conductor cable running between the Power Distribution Board’s 12V and GND rails to a 5V relay board’s NO and COM pins set to low trigger (COM-15093). The removal of a 5V digital input coming from the LabJack T7 during the transition from the 12 night-hours to the 12 day-hours completes the circuit and turns on a constant lux LED strip to simulate daylight conditions. The NO pin of the relay board and the positive lead of the 9 cm (3.5 inch) long LED strip are connected via a female-male barrel jack junction. The LabJack T7 provides the power relay board with 5V of input during the 12 night-hours and removes it during the 12 day-hours. In the absence of that 5V input, the power relay board supplies 12V of power to the visible light LED strip.

### Arduino-Motor Shield Construct

This consists of an Adafruit Motor Shield Assembly (1438 v.2) mounted on an Arduino Mega 2560 (Product number A000067). DC power is split as described above, with 12V for the Arduino and 9V for the stepper motor output of the mounted Motor Shield. The shield also uses 5V from the Arduino to power its logic on-board.

Arduino Mega 2560 links beam break inputs at the nose poke ports to the water or food dispensing outputs to be mediated by the Motor Shield (opening valves or powering motor rotations). The Arduino code can be calibrated to control the water dispense drops of water via modification of the time the solenoid valve remains open. Also a “timeout” function in the code ensures dispenses cannot happen with an interval of less than 3 seconds regardless of nose pokes in the 3 second refractory period. Regardless of the timeout configuration, Arduino records all beam break events and relays the information to the LabJack T7 via an IDC parallel cable. All dispense events are also recorded to LabJack.

In the Motor Shield Assembly, the white, red and green wires of 6 female 3-pin connectors (CAB-14575) are soldered onto their corresponding inputs in the digital I/O rail, power rail, and ground rail of an Adafruit Motor Shield V2, respectively. Four (4) of these connectors interface with the 4 different Beambreak Assemblies (sucrose and regular water, fatty and regular food). Soldering the connectors to the Motor Shield Assembly instead of the Arduino allows our system to remain modular, and enables the Arduino to easily be reprogrammed, tested, and swapped out if needed. The Motor Shield receives a 5V digital input in the event of a beam break at the dispensing ports, and according to I/O relationships encoded by the Arduino, it either uses 9V to energize one of the coils in the appropriate stepper motor or sends a 5V digital logic input to the N-channel enhancing type MOSFET of the solenoid driving circuit. This input allows current flow through the appropriate solenoid valve for a user-defined period, resulting in water dispensing. An IDC parallel cable transfers digital information encoding all beam break and dispensing events from the Arduino to the LabJack T7, which also directly receives analog data encoding running wheel revolutions.

### Food Dispensing and Water Dispensing

Dispensing Structure: The 3D-printed structure was made using Prusa MK4 with PLA/PETG resin (Overture) or Formlabs2 3D Printer, rigid resin (FLRGWH01) and is composed of one main body holding two stepper motors to rotate each of two food hopper wheels and two food holders above the hoppers, two chutes for hopped food to descend to the mouse, and a mouse interface printout. The mouse interface includes four nose poke beambreaks, two receptacle catches for each of the food chutes, and water dispenser needles for water dispense. This design was based off of an open-source design from https://open-ephys.atlassian.net/wiki/spaces/OEW/pages/79069188/Food+Pellet+Dispenser. This was followed by our own modifications after converting the .stl files into editable SolidWorks files.

The food dispensing works via a chute that delivers food from an elevated receptacle when the mouse pokes its nose in the beambreak of the port. A servo motor turns to drop the food down the chute to a food presentation portion of the mouse port – a few millimeters below the food beambreak zone. The upper dispenser under the receptacle has slits to prevent debris accumulation (Figures 2Ai). Two water ports are similar, dispensing either regular or 1% sucrose water (Figure 2Aii, left). These work via a nose poke beambreak leading to the opening of a solenoid to let water flow down a tube from a suspended syringe.

Beambreak Assembly: Our system employs four Beambreak Assemblies, one at each dispensing port, consisting of an optical switch photointerruptor relying on infrared light (Mouser Electronics 474-BOB-09322), a male 3-pin connector that interfaces with the female connectors of the Motor Shield Assembly, and a 220Ω current-limiter resistor to protect the infrared LED. The Motor Shield receives a 5V digital input in the event of a beam break, and according to I/O relationships encoded by the Arduino, it either uses 9V to activate one of the coils in the appropriate stepper motor or sends a 5V digital logic input to the N-channel enhancing type MOSFET of the solenoid driving circuit.

Food: Regular (4 mm diameter) pellets and 15% purified medium-fat pellets (Bio-Serv Dustless Precision Pellets F0165 and F0021, respectively) are provided *ad libitum* for the mouse via the two separate food dispensing units located inside the mouse cage. When a nose-poke is detected by the Beambreak Assembly at one of the food-dispensing ports and the digital signal is processed as outlined in the Arduino-Motor Shield Construct, the Motor Shield ends up supplying 9V, 500 mA to a corresponding bipolar, 2-coil stepper motor (Pololu-1208, SY35ST28-0504A) located in the 3D printed Dispensing Structure. The stepper motor has a 1.8° step angle, offers a holding torque of 0.09 Nm and is connected to the Motor Shield via four wires, two for each motor coil. The stepper motor shaft is lodged into the pellet distributor disk and is lined with eight, 0.635 cm (1/4 inch)-diameter, evenly spaced holes that fit a single pellet at a time. To prevent clogging of the system with pellet debris, the receptacle base is lined with rectangular slits and the direction of motor rotation is reversed once every 5 steps via Arduino code.

Water: Filtered water and 1% sucrose water are provided *ad libitum* for the mouse via the two separate water dispensing ports inside the cage. Two 10 mL syringe shafts (Bio-Rad: M10LLASSM), one filled with 1% sucrose water and the other with distilled regular water, are connected to the input port of the corresponding solenoid valve via a 76 cm-long, 1.6-mm ID Tygon water tube (Bio-Rad: 7318215) fitted with a female lure lock adaptor on the syringe shaft end. The open end of a similar 10 inch-long tube connects to the output port of the solenoid valve. The other end arrives at the water dispensing port and is fitted with a male lure lock adaptor and a 20-G x 1.5-inch needle (Medline SYR100207) with the bevel removed and tip dulled to prevent any injuries to the mouse. The Motor Shield supplies 5V of constant power to the Water Solenoid Valve Driving Circuit. The circuit also includes a Zener diode (1N4757A) for overvoltage protection, a rectifier diode (14005-T) for forward biasing, and a 10 kΩ (CF14JT10K0CT-ND) pulldown resistor.

### Running Wheel System

The system uses a commercially available mouse running wheel (Bio-Serv Fast Trac and Mouse Igloo K3250) made of autoclavable amber polycarbonate transparent to infrared light, preventing it from blocking our infrared camera. The interface between the igloo and the trac was modified to accommodate a stainless steel, 1.07 cm (0.42 inch)-diameter shaft coupler (ServoCity: 625152) fitted with a wooden 4.7 mm (3/16-inch) diameter x 3.56 cm long wooden Dowel rod that lodges into the Fast Trac on one end and a MA3 Miniature Absolute Magnetic Shaft Encoder (US Digital MA3-A10-125-B) on the other. This shaft encoder generates a 10-bit analog voltage output proportional to the absolute shaft position. This analog output voltage is recorded by our LabJack by means of a 4-conductor shielded round cable with a 3-pin micro mating connector on one end (CA-MIC3-SH-NC unterminated with shielded cable, US Digital). Here, the voltage at every half rotation is recorded, timestamped, and continuously appended to a .csv file containing all analog data.

### Infrared Video Recording System

Our system makes use of a camera (Basler, acA1300-200 um) with infrared-pass filter (FRMS208 Nir Bandpass filter, Machine Vision Store) that delivers 20 frames/second at 1.3 MP resolution. The camera connects to a desktop computer via USB and produces an .mp4 video file every 4 hours. For video imaging, animal and homecage are illuminated an infrared LED backlight that remains perpetually lit behind a translucent plexiglass panel that diffuses the light, allowing for even illumination at the infrared camera. The wavelength of the infrared light falls below the detection threshold for mice, allowing for continuous recording, even in dark hours. Visible light is modulated daily, while this infrared light remains on constantly. BandiCam software is used to record frames reliably and also to start and stop acquisition every 4 hours. Timestamped video files are then analyzed with both HomeCageScan (CleverSys) and MATLAB code to extract video-based actigraphy.

### Software Ecosystem

Arduino Integrated Development Environment (IDE): contains code to control the Arduino-Motor Shield Construct. Its functions determine input-output relationships in the Motor Shield underlying food and water dispensing. This allows for testing and manipulating the operation of the stepper motors and solenoids, diagnosing their failure causes, and troubleshooting potential issues with functions that can, among other things, unjam clogged dispensing mechanisms.

phoBehavioralBoxLabjackController: This custom software written in C++ collects running wheel revolutions, beam break and dispense event data, and controls the toggle of the warm yellow LED strip via the LabJack T7. The program produces output in the form of two .csv files, one for digital data and the other for analog data.

LabJack custom software: Writes a new line to the digital (food/water nose poke/dispense) or analog .csv file whenever a voltage change is detected. In each line, the full state of all digital or analog pins is written using booleans (digital) or 0-5 voltage (analog) alongside POSIX timestamps. The LabJack T7 uses a rate of 50 kHz.

### Digital Clock Synchronization

Each of our embedded devices operates on its own internal clock. Desktop computers come with an accurate real time clock built-in, which is frequently synchronized with the NIST atomic clock. This allows POSIX timestamping of data from each stream to enable time-of-day analyses.

## QUANTIFICATION AND STATISTICAL ANALYSIS

All analysis (food/water/running wheel/video) was cleaned and organized as described below and data were then converted to zeitgeber time (ZT) with hour ZT0 at lights on and hour ZT12 at lights off. The default value for each analysis was hours; however, for continuous actions such as wheel running, minute data was also extracted for duration measurements.

Food and Water: Using Matlab (Mathworks, 2024b) we binned (by hour) digital data for beambreak and dispenses from the RShiny server as separate CSVs for each animal across the experimental period. First, new rows were added for hours which had no recorded beambreaks/dispenses and a count of 0 assigned. Across all DHCs, hours with more than 500 beambreaks were noted and investigated for dispenser clogging or leaking. If the video for that hour showed dispenser malfunctions or animal activity that did not match beambreak/dispense levels (mouse was sleeping or did not interact with the feeder), both the beambreak and dispense value for that hour were replaced with ‘Not a number’ (NaN) and excluded from all analyses. For analysis by week across animals, for each animal, hourly beambreaks were summed across each week of the experiment for each DHC. For analysis by day across animals, hourly beambreaks and dispenses were summed across each day for each animal then averaged across all animals for each day. For analysis by hour across animals, average was calculated across all available hourly beambreaks and dispenses for all animals for each hour of the day. All data was graphed using GraphPad Prism (GraphPad v.10).

Place Preference: Beambreaks for both food and water types were summed for each day of the experiment across all animals. Values for the daily beambreak sum at each food and water position were parsed from the daily beambreak type values, assigning each daily sum for a type of food or water to the proper position as marked by the researchers in their daily box check. Next, daily differences were calculated across position and type sums, always calculated as (position 2 - position 1) or (fatty/sucrose – regular). For food and water type data, differences were placed in one of two position groups depending on the position of sucrose water or fatty food for that day. For position data, differences were placed in groups depending on what type of food or water occupied position 1 for that day.

Running Wheel: We downloaded minute- and hourly-binned analog data for distance run from the RShiny server as separate CSVs for each animal across the experimental period. Per-minute cumulative distance across the experiment was calculated for each DHC by replacing the value of each cell in the csv with the sum of itself and all values before it. These data were then plotted for each DHC and investigated for non-physiological activity (large spikes or sustained activity during the light period). Abnormal periods were compared to available video data, and animal activity that did not match reported distance values were replaced with ‘NaN’ and excluded from analysis in hour chunks from both binned files for that DHC.

Cleaned data were then converted to ZT in both hour and minute bin files. Per-minute cumulative distance across the experiment was recalculated for each DHC using the same methods as previously described. Similarly to the digital data, new rows were added in the hour binned file for hours which had no recorded wheel running, and a distance of 0 was assigned to that hour. Linear regression was calculated from every hourly binned data point available for the acclimation and baseline periods across each DHC. For analysis done by day across animals, hourly wheel run distances were summed across each day of the experiment for each DHC. For analysis of time per hour spent running, the minute bin entries were counted across each hour of the experiment for each DHC.

To generate vectors, time in hours was first converted to radians to map the 24-hour cycle onto a circle. The vector’s components were calculated by summing the weighted horizontal (x) and vertical (y) contributions of distance or time averages across all hours, using cosine and sine functions, respectively. The mean direction of distance or time was then derived from these components using the arctangent function and converted back to hours. The vector’s magnitude, reflecting the concentration of distance or time around the average hour, was added using vector addition to yield a mean resultant vector, yielding a vector with mean direction and magnitude. This process was performed on each DHC for hourly distance, average distance, and time values across all animals.

### Video and Behavior--HomeCageScan

HomeCageScan (HCS) was used for behavioral video analysis from CleverSys Inc (Reston, VA). Videos from the same mouse consecutive in time (usually a week of videos between cage changes) are nominated for batch HCS processing as a “session”-- or indices of the locations of a desired .mp4 video file. Text log files of processing steps for the selected video are saved to assist in later data processing. HCS session initiation requires the outlining of a region of interest to indicate the “cage”, or space wherein the mice will behave, and a static background image to compare movement to. Feeding and drinking zones are also indicated manually as polygons. As video is processed, the software will index for the requested video, apply the virtual cage, and automatically regenerate background images if the program loses track of the animal’s location. Video is processed at maximum speed, 3000 frames per second. Behavioral data are recorded in duration of frames and saved to a .csv file, which is then processed to extract the periods of bouts of certain behaviors. Custom Python software takes in both the finished .csv files, and the text log file containing the video indices and inserts the frame-based POSIX time of each behavior on the .csv.

To create the 14 primary categories analyzed, behaviors in the scored .csvs were combined by similar behavior type (walk left/right/slowly, eat zone 1/2, drink zone 1/2, etc.) across all animals. Fifteen behaviors comprising approximately 93% of animal activity were retained, and the remaining behaviors were grouped in with the existing ‘Other’ category defined by HCS.

Time spent in each behavior per hour was calculated across all available days of data by first iterating over each annotated file, accumulating total durations (converting frames to second) per behavior per hour of the day. The total duration of each behavior per hour was then divided by the number of times that hour occurred in the dataset, returning the time spent in each behavior per hour. Vectors for hourly behavioral activity were calculated using the same methodology as described for the running wheel.

The example 24-hour and two 12-hour behavior breakdown bar charts were calculated by summing the frames spent in each behavior then converted to seconds based on a consistent frame rate of 30 fps. Total seconds spent in each behavior was divided by 86400 seconds (one day) to determine the percentage of time spent in that behavior.

---

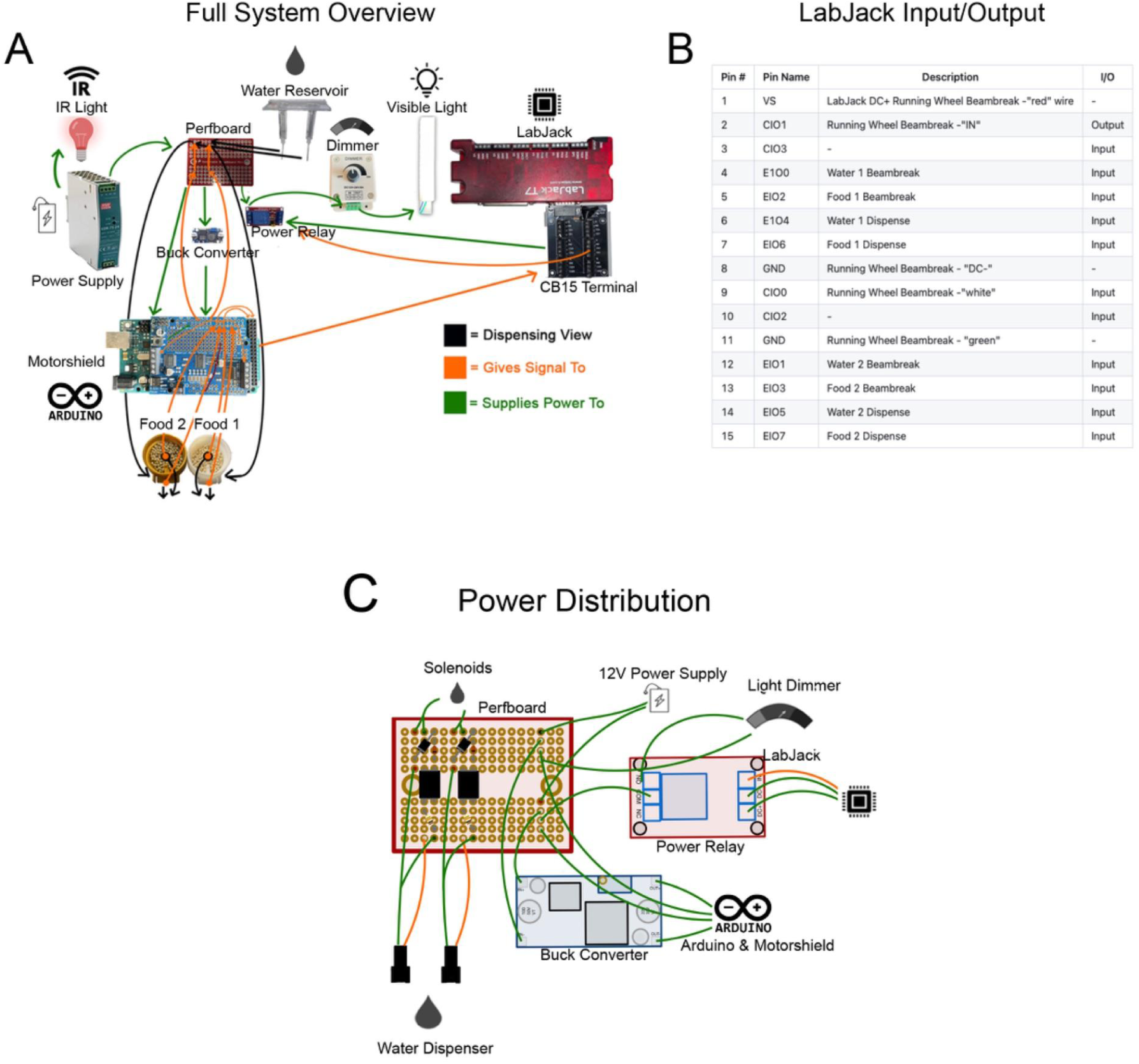

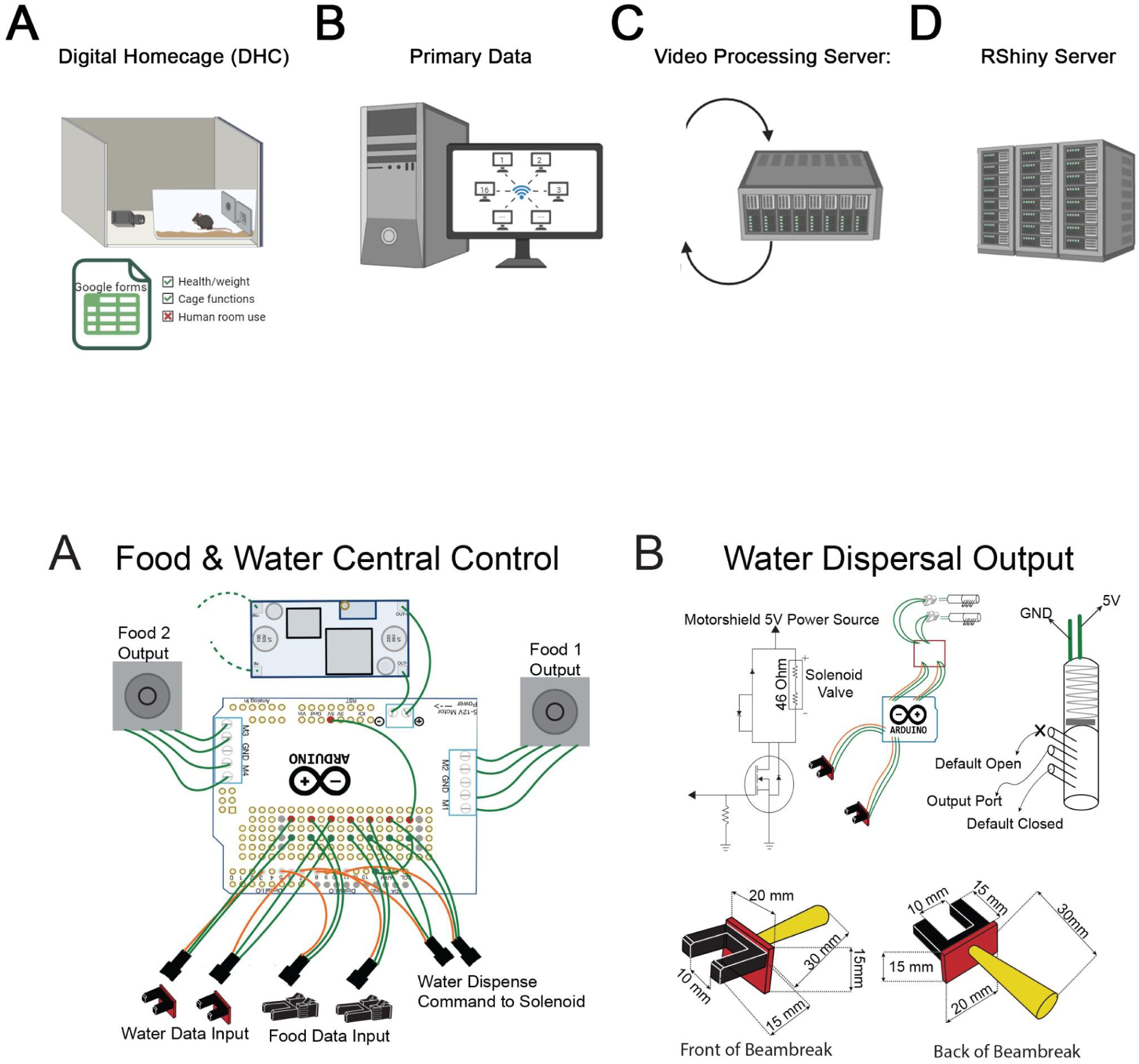

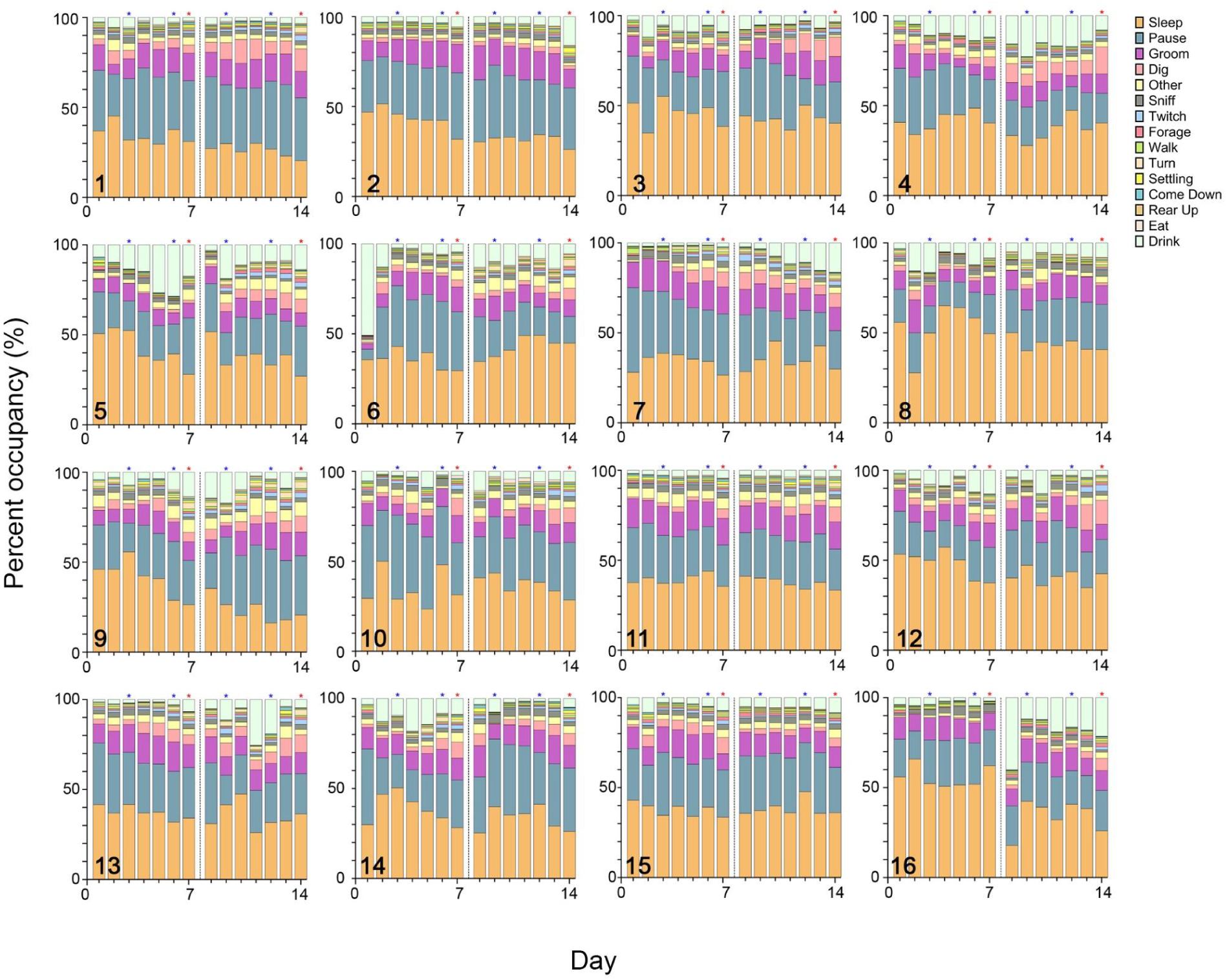

